# FLASHDeconv: Ultrafast, high-quality feature deconvolution for top-down proteomics

**DOI:** 10.1101/714915

**Authors:** Kyowon Jeong, Jihyung Kim, Manasi Gaikwad, Siti Nurul Hidayah, Laura Heikaus, Hartmut Schlüter, Oliver Kohlbacher

## Abstract

Feature deconvolution, the determination of intact proteoform masses, is crucial for top-down proteomics, but currently suffers from long runtimes and quality issues. We present FLASHDeconv, an algorithm based on a simple transformation of mass spectra, which turns deconvolution into the search for constant patterns thus greatly accelerating the process. We show higher deconvolution quality and two to three orders of magnitude faster execution speed than existing approaches.

---

Top-down (TD) proteomics has gained a lot of momentum for in-depth protein characterization and protein species analytics^1–7^. In contrast to bottom-up (BU) proteomics, where proteins are enzymatically digested and actually peptides are analyzed, the TD approach allows for the analysis of intact proteoforms (protein species arising from the same gene product via splice variants, genomic variation, post-translational modifications, degradation, etc.)^6, 8^. Due to the high mass of the analytes, however, in mass spectrometry (MS)-based TD approach (TD-MS), a single protein species can result in multiple MS features with different charge states and isotopic envelopes, making the signal structure highly complex and, in turn, the accurate determination of proteoform masses challenging^9^. Feature deconvolution (i.e., the determination of intact proteoform masses) is thus an essential step for TD data analysis^10, 11^.

Current MS instrumentation and experimental protocols enable more complex analyses using TD proteomics, but in turn yield equally complex and large data sets^1, 2, 6, 7, 10, 12^. Hence, both quality and runtime of deconvolution algorithms have become an issue. Current deconvolution algorithms typically have long processing times of hours or days for complex data sets. Quality is another issue of these approaches. Mass artifacts and limitations with respect to mass or charge ranges reduce the methods’ applicability^10, 11, 13^.

We present FLASHDeconv, an algorithm for high-quality TD deconvolution two to three orders of magnitude faster than existing tools. The major speed-up of FLASHDeconv is achieved by very fast decharging (i.e., assigning charges to peaks) in the spectral deconvolution step (Fig. 1a). It is based on a simple transformation of mass spectra that transforms peak positions (m/z) to log m/z space. The variable (mass-dependent) patterns caused by different charge states of the same protein are turned into a mass-independent universal pattern by this transformation. Finding the occurrences of the universal pattern in a transformed mass spectrum can be done very efficiently by a single convolution calculation. FLASHDeconv also implements a deisotoping algorithm that works well for both isotopically resolved and unresolved regions and simple yet effective filtering/scoring schemes to reduce false positives (see online methods).

**Figure 1.**
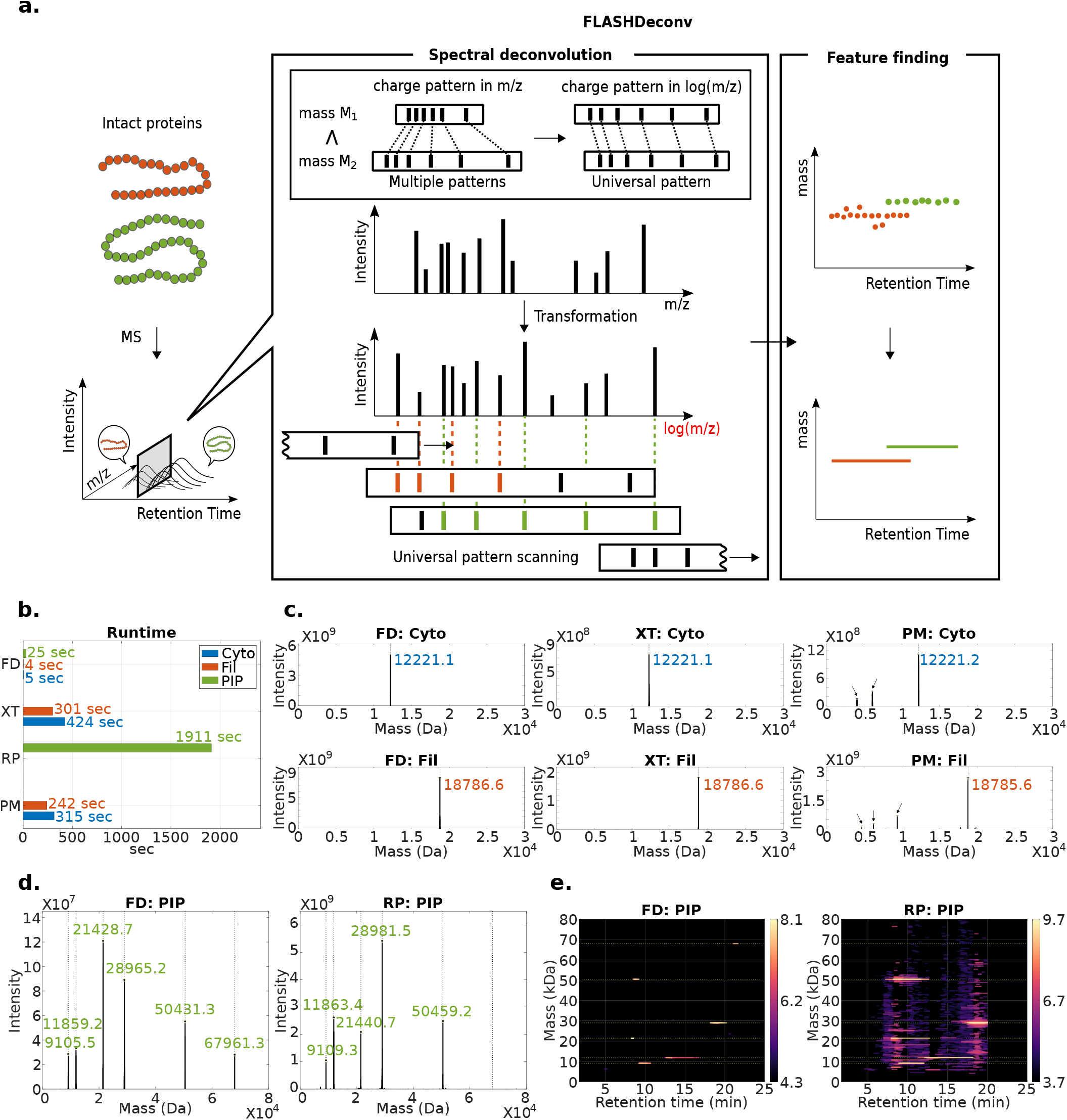
FLASHDeconv algorithm and analysis of simple data sets. **a**. FLASHDeconv consists of two separate processing steps: spectral deconvolution and feature finding. Spectral deconvolution finds monoisotopic masses from each TD-MS spectrum. The upper box illustrates the key idea for rapid deconvolution: if each peak position is transformed into log of its neutral m/z value, the charge pattern (the position pattern caused by different charge states of the same protein) in the transformed spectrum becomes independent of protein masses and hence is universal. Thus, decharging can be done by a single sliding this universal pattern along the transformed spectrum (see online methods) to identify matching peaks. Feature finding then reduces to finding agreeing deconvolved masses within given mass tolerance along the retention time (RT) direction (done by a mass trace detection algorithm^19^). **b**. Run time comparison for simple data sets (Cyto, Fil, and PIP; see main text for abbreviations). FD stands for FLASHDeconv; XT for Xtract; RP for ReSpect; and PM for Promex. FD was up to 90 times faster than the other methods. **c**. Deconvolved spectra generated by FD, XT, and PM for isotopically resolved pure protein data sets(Cyto - upper panel; Fil - lower panel). Dotted black lines indicate the expected exact masses (12,223.21 Da and 18,802.8 Da), and the colored digits specify the reported masses within 3 Da from exact input monoisotopic masses. For intensity estimates, MaxIntensity column was used for FD, Sum_Intensity for XT, and Abundance for PM. All three methods found the expected masses, but PM reported additional high-intensity low-harmonic mass artifacts (e.g., ½ and ⅓ input masses; marked with arrows). **d**. Deconvolved spectra from FD and RP for the PIP (six standard intact protein mix) data set (isotopically unresolved). The colored digits specify the reported masses within 3 Da from input monoisotopic (for FD) or average (for RP) masses. For intensity estimation, Sum_Intensity was used for RP. FD found all six input protein masses. RP found five. The expected exact monoisotopic masses are 9,105.3, 11,858.0, 21,429.8, 28,963.7, 50,429.8, and 67,959.4 Da. The expected average masses are 9,111.5, 11,865.5, 21,442.6, 28,981.3, 50,459.7, and 68,001.2 Da. **e**. Identified features mapped on the RT-mass plane (feature maps) from FD and RP for the PIP data set. Except for the five expected masses, RP also reported thousands masses, out of which 43% were identified as mass artifacts or isotopologues (see online methods for the artifact detection method). See also Supplementary Fig. 1-2 for more results from the replicates.

To evaluate its performance, we benchmarked FLASHDeconv with TD samples of varying complexity (three low complexity samples, Fig. 1b-e; and one highly complex sample, Fig. 2). The three low-complexity samples consist of individual proteins (Cyto - Bovine Cytochrome C, Fil - Filgrastim) and a standard protein mix (PIP - Pierce Intact Protein Standard Mix; see online methods for details). Cyto, Fil, and PIP data sets were acquired on a Quadrupole-Orbitrap at a resolution of 70,000 (Cyto and Fil data sets) and 17,500 (PIP). Due to their low masses (12,223.21 Da for Cytochrome C and 18,802.8 Da for Filgrastim) and high resolution, the spectra in the Cyto and Fil data sets contain isotopically resolved peaks. The PIP data set was acquired at a lower resolution of 17,500 to assess the deconvolution quality for unresolved isotope patterns. For the complex data set, we used a murine myoblast sample (unfractionated and generated by Q-Exactive HF at a resolution of 120,000) published previously^14^. This data set contains both isotopically resolved and unresolved peaks due to the wide mass range.

**Figure 2.**
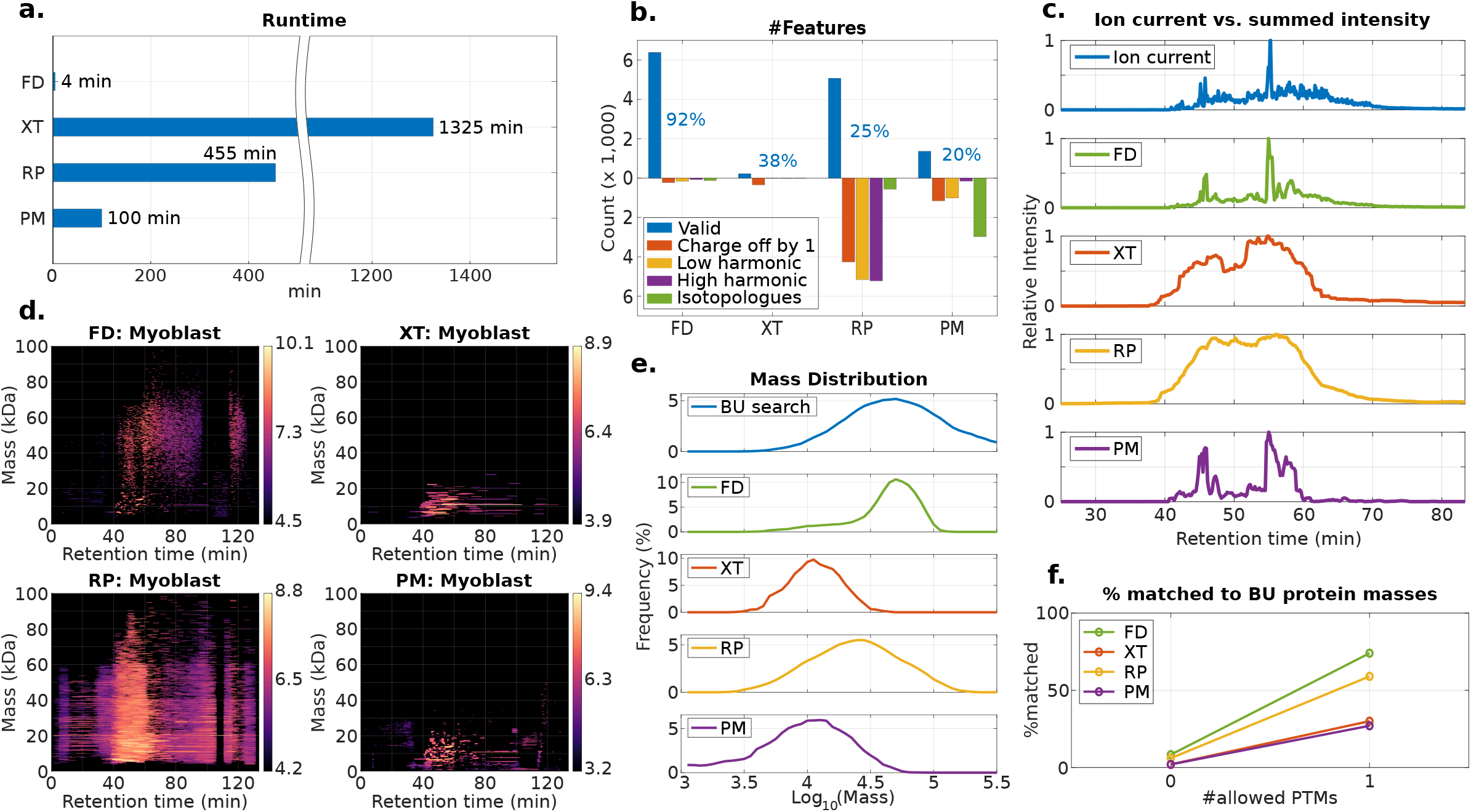
Analysis of complex murine myoblast data set. **a**. Runtime comparison for the murine myoblast data set. FD was drastically faster (25-330 times) than the other methods. **b**. The numbers of valid features (i.e., not detected as artifact) and detected mass artifacts/isotopologues from each tool (see online methods for the artifact detection method). The blue digits show the portion of the valid features. FD found almost 7,000 features, and only 8% of them were artifacts. The other tools generated significantly higher portion of artifacts (60-80%). **c**. For each tool, feature intensities were summed across all masses and plotted along RT direction. For reference, total ion current (TIC) chromatogram was also drawn on top. FD and the TIC showed almost perfect fit, demonstrating FD features cover most of eluted ions. **d**. The feature maps for each tool. For the 0-20 kDa region, all tools produced intense features (see Supplementary Fig. 3 for rounded monoisotopic mass overlaps in low mass region). XT and PM reported only few features for 20-100 kDa region, because most of the corresponding peaks would be isotopically unresolved. **e**. The mass distribution of each tool as compared to the protein mass distribution obtained from bottom-up (BU) search (of the same myoblast sample). The distribution from FD is similar to the BU identified protein mass distribution. **f**. The portion of the feature masses matched to the BU identified protein masses for each tool (see Supplementary Table S1). FD exhibits a much higher consistency with BU results than other tools.

For comparison, Xtract^15^ (in Thermo BioPharma Finder 3.1), ReSpect (Positive Probability Ltd, in Thermo BioPharma Finder 3.1), and Promex^16^ (version 1.0.7017) were tested against FLASHDeconv for selected data sets: Xtract and Promex for Cyto, Fil, and myoblast; ReSpect for PIP and myoblast. Detailed sample generation procedure and tool parameters are found in the online methods.

Fig. 1b-e shows the results for the simple protein data sets (see Supplementary Data 1 for the full list of features). In Fig. 1b, it is shown that FLASHDeconv is 60-90 times faster than comparable tools. Per spectrum, the deconvolution took less than 6 ms (excluding spectrum loading time). Fig. 1c shows the deconvoluted spectra (deconvoluted masses and their intensities, merged for all RT range) for the isotopically resolved Cyto and Fil data sets. All three tools found a signal around the input protein masses within 3 Da mass error. However, Promex reported additional harmonic masses (e.g., ½ and ⅓ of the input masses) for both data sets.

In the isotopically unresolved PIP data set, FLASHDeconv found all six input masses within 3 Da mass error (ranging from 9 to 70 kDa; Fig. 1d-e). ReSpect found five input masses, excluding the heaviest one. The features from each tool were also mapped to the RT-mass plane in Fig. 1e (feature map). The feature map from ReSpect shows that ReSpect reports thousands of features in addition to those directly relatable to the input masses. However, when we examined the mass and RT relations of the features, we found that at least 43% of them were classified as mass artifacts or isotopologues (see online methods). The results on the low-complexity samples demonstrate that FLASHDeconv accurately identifies the proteoform masses with high specificity and sensitivity for both isotopically resolved and unresolved data sets. Supplementary Fig. 1-2 provides the same information as Fig. 1b-e for replicate data sets, where the almost identical results were reproduced (see Supplementary Data 2-3 for the complete feature lists).

The advantage of FLASHDeconv is even more pronounced on the complex data set (see Supplementary Data 4 for feature lists). Fig. 2a shows the run time comparison for the myoblast data set. FLASHDeconv was significantly faster (up to 330 times faster than Xtract) than the other tools; it took only four minutes to process the whole data set (approx. 60 ms/spectrum).

We found that the numbers of reported features differed significantly from tool to tool (from around 600 for Xtract to about 20,000 for ReSpect). We then examined the masses and RT relations among the features as above (see online methods) to assess whether mass artifacts or isotopologues contribute to this large feature count variation between tools (Fig. 2b). For FLASHDeconv, only 8% of the detected features actually turned out to be artifacts or isotopologues, whereas that number reached almost 80% for some of the other tools. Excluding the detected artifacts, the largest number of features (~6,000) were reported by FLASHDeconv, demonstrating its high sensitivity and specificity. To see this in a different perspective, we compared the total ion current (TIC) chromatogram (from the raw spectrum file) with the intensities summed over masses (from the reported features) in Fig. 2c. FLASHDeconv showed almost perfect fit to the TIC. All other methods show a much lower agreement with the TIC. Good agreement between the TIC and the summed-up feature signal basically indicates that a large portion of the recorded signal could be assigned to the detected features. Poor agreement can occur when parts of the signal are not detected (causing sensitivity drop) and/or false positive features are erroneously included (causing specificity drop).

The feature maps in Fig. 2d and the feature mass distributions in Fig. 2e together show FLASHDeconv features are centered between 40-60 kDa while the others between 0-20 kDa or 0-40 kDa (see Supplementary Fig. 3 for the rounded monoisotopic mass overlap between tools in low mass region). Not surprisingly, Xtract and Promex exclusively reported masses lower than 20 kDa, as most masses larger than 20 kDa would be represented by isotopically unresolved peaks (for which Xtract and Promex are not optimized). Next, we compared our feature masses with the protein masses identified by BU searches for the same sample (also reported by L. V. Schaffer et al., 2018^14^). The distribution of these BU-identified protein masses (Fig. 2e top panel) was similar to the FLASHDeconv feature mass distribution, suggesting that the high masses reported by FLASHDeconv are indeed present in the sample. For more rigorous evidence, lastly we matched our feature masses against the BU identified protein masses (Fig. 2f; see online methods for matching method). About 74% of the feature masses were matched when a single modification was allowed, confirming most of our feature masses are likely to be genuine proteoform masses. The other tools showed substantially lower agreement than FLASHDeconv (see Supplementary Table 1 and 2).

In summary, FLASHDeconv can carry out high-quality deconvolution for diverse types of (simple and complex; and isotopically resolved and unresolved) TD-MS data sets at unprecedented speed and quality. With its high speed and versatility, FLASHDeconv could significantly increase the potential of various TD analyses including native-MS and TD-BU integrated studies^4, 17^. Lastly, the very fast processing times for spectral deconvolution could be used for real-time mass deconvolution on MS instruments, enabling smarter and more efficient MS/MS fragmentation methods for TD-MS than are currently being used.

## Supporting information

Supplementary Data 1 (Cyto, Fil, and PIP)

Supplementary Data 2 (Cyto, Fil, and PIP rep. 1)

Supplementary Data 3 (Cyto, Fil, and PIP rep. 2)

Supplementary Data 4 (myoblast)

## Code availability

FLASHDeconv is implemented in C++ as a part of OpenMS^18^ and available as platform independent open-source software under a BSD three-clause license at https://OpenMS.org/FLASHDeconv.

## Data availability

The raw files for Cyto, Fil, and PIP triplicate data sets have been uploaded to MassIVE (https://massive.ucsd.edu) and are available under accession number MSV000084001 or under the digital object identifier https://doi.org/10.25345/C59D26.

## Acknowledgements

We are very thankful to Lloyd M. Smith for generously making murine TD&BU data sets publicly available. We thank to CinnaGen Co. for sharing Filgrastim protein and are grateful to Christoph Henrich, Bernard Delanghe, David Horn, and Jennifer Sutton from Thermo Fisher Scientific for insightful discussions. K.J. is grateful to Jong-Seo Kim and Jong-Eun Park for their advice on the paper. J.K., M.G., S.N.H., L.H., H.S., and O.K. acknowledge funding from the Horizon 2020 Marie Sklodowska-Curie Action ITN 2017 of the European Commission (grant 765502-A4B). O.K. acknowledges funding from BMBF (de.NBI, 031A535A).

## Contributions

O.K. conceived the idea of the fast deconvolution algorithm for TD-MS data sets. K.J. and J.K. developed and implemented FLASHDeconv algorithm. M.G., S.N.H., L.H., and H.S. designed the experiment and performed sample preparation and data acquisition. O.K. and H.S. led the project and provided resources. K.J. and J.K. wrote the manuscript with input from all authors. All authors commented on and approved the paper.

## Online methods

### FLASHDeconv algorithm

FLASHDeconv consists of two sub-algorithms, spectral deconvolution and feature finding. FLASHDeconv performs fast mass deconvolution of each spectrum, and subsequent mass trace detection along retention time in the deconvolved spectra.

### Spectral deconvolution algorithm

#### Decharging and harmonic artifact removal

FLASHDeconv reads mass spectra in HUPO-PSI-compliant mzML format^20^ and then deconvolves each of these spectra individually. For this study, we used ProteoWizard^21^ msconvert tool (without peak picking or filtration option) to convert raw spectrum files to mzML files. Goal of the decharging is to identify all peaks caused by the same protein mass, where the peaks are differing only with respect to their charge states. No initial peak filtration is applied by default since even low-intensity peaks can be important evidence for low-abundance features. Finding these groups of related peaks (peak groups) is achieved by searching for a mass-invariant charge pattern in log-transformed m/z space (see main text, Fig. 1 for the concept).

For each input spectrum ***S***, FLASHDeconv computes a transformed spectrum ***S\**** by subtracting the mass of the charge carrier (e.g., in positive mode, a proton mass) and logarithmizing (natural base) the resulting value for each peak. We denote the mass of the charge carrier by ***q***. As m/z of a peak of charge ***c*** from mass ***m*** is given by (***m*** + ***cq***)/***c*** = ***m*** /***c*** + ***q***, the transformed peak position is at log(***m*** /***c***) = log(***m***) − log(***c***). Thus, for a series of peaks ***p***_***1***_, ***p***_***2***_,…, ***p***_***n***_ from the same mass ***m*** with charges ***c***_***1***_, ***c***_***2***_,…, ***c***_***n***_, the transformed peak locations are given by log(***m***) − log(***c***_***1***_), log(***m***) − log(***c***_***2***_),…, log(***m***) − log(***c***_***n***_). We can thus define a universal charge pattern ***U*** := (−log(***c***_***min***_), −log(***c***_***min***_ + 1),…,−log(***c***_***max***_)) for a given charge range [***c***_***min***_, ***c***_***max***_] (two and 100 by default). Then, the peaks in the peak group of mass ***m*** should partially match (within tolerance) the universal charge pattern ***U*** positioned at ***log*** (***m***). Similar to the universal charge pattern ***U***, we define the harmonic charge patterns ***H***_***r***_ for ***r*** =½, ⅓, ⅖ as ***H***_***r***_ := (−log(***c***_***min***_ + ***r***), −log(***c***_***min***_ + 1 + ***r***),…,−log(***c***_***max***_ + ***r***)). Harmonic charge patterns are used to filter out low harmonic artifacts (e.g., the masses having ½, ⅓, ¼, ⅕, or ⅖ times the true mass). Further filtration of (both high and low) harmonic artifacts are performed after peak groups are defined below.

FLASHDeconv slides ***U*** and ***H***_***r***_ along the transformed spectrum ***S^*^*** to find suitable matching peak groups. Let ***U*** (***x***) denote the pattern ***U*** at the position ***x*** in ***S^*^*** (likewise, ***H***_***r***_(***x***)). Note that ***U*** (***x***) is matched by the peaks from mass ***exp***(***x***). Two peaks ***p*** and ***p’*** are called comparable if their intensity ratio is between ¼ and 4. We call two peaks ***p*** and ***p’*** are well-aligned if they are comparable and they match to any of two consecutive elements of ***U*** (***x***). A peak ***p***_***h***_ is a low harmonic peak with respect to well-aligned ***p*** and ***p’*** if i) ***p***_***h***_ is located between ***p*** and ***p’***, ii) ***p***_***h***_ is comparable with ***p*** and ***p’***, and iii) ***p***_***h***_ matches to any element of ***H***_***r***_ (***x***). Then, two well-aligned peaks ***p*** and ***p’*** are further called signal peaks if they have no low harmonic peak.

While sliding a pattern along the transformed spectrum, FLASHDeconv stops at the location ***x*** where three consecutive signal peaks ***p***_***1***_, ***p_2_***, ***p***_***3***_ are detected, that is, (***p***_***1***_, ***p***_***2***_), and (***p***_***2***_, ***p***_***3***_) are pairwise signal peaks. Then it recruits all peaks matched to ***U*** (***x***), forming a peak group of mass ***exp*** (***x***). A peak matched to an element −log(***c***) in ***U*** (***x***) now has the assigned charge of ***c***.

Even though the method described above efficiently collects peak groups and assigns charges to their peaks, still multiple peak groups represent either low or high harmonic artifacts. Also, quite often a single peak has multiple assigned charges. This happens mainly when a single peak has membership in multiple peak groups, often those representing harmonic artifacts. To resolve this, we assign an intensity score (described below) to each peak group. Briefly, the intensity score measures the total signal peak intensity subtracted by the total harmonic (both high and low) peak intensity. Then, for each peak, we collect all peak groups containing the peak. The peak group of the highest intensity score is allowed to retain the peak. The peak charge is updated accordingly. At this point, the set of peak groups is called the candidate peak group set. The mass of a candidate peak group is determined by the most intense peak mass therein.

To define the intensity score of a peak group, we detect possible harmonic peaks in the peak group. The low harmonic peaks are already detected when finding signal peaks. To detect high harmonic peaks, first, we sort the peaks in ascending order of their assigned charges. Denote the ***i***-th peak in the sorted peak list as ***p***_***i***_. For each peak ***p***_***i***_, we take the peaks ***p***_***i*−*1***_ and ***p***_***i*+*1***_. The peak ***p***_***i***_ is called a high harmonic peak if ***p***_***i***_ is comparable but is not well-aligned with ***p***_***i*−*1***_ and ***p***_*i*+*1*_. For all collected signal, low harmonic, and high harmonic peaks in the peak group, the intensity score of a peak group is defined as its total signal peak intensity subtracted by its total low and high harmonic peak intensities.

The number of candidate peak groups is usually less than 2,000 even for complex spectra. Thus complicated processing of the candidate peak groups from this point on does not slow down FLASHDeconv significantly.

#### Peak group clustering for isotope envelopes (isotopically resolved case)

Each peak group has its mass, but some of them are from the isotopologues of the same protein. Moreover, some of the peaks in ***S\**** may not have been selected for any peak group due to their low abundance even though they are isotopologues of existing peak groups. Thus, after defining candidate peak groups, we cluster peak groups and recruit additional peaks from plausible protein isotopologues.

This processing is done in a greedy fashion from the peak group with the smallest mass to the one with the largest mass. Let ***m***_***iso***_ be the mass difference between ^13^C and ^12^C. Suppose we are processing a peak group ***P*** of mass ***m***. First, we examine if there are peak groups **P’** of mass ***m*** − ***m***_***iso***_ or ***m*** + ***m***_***iso***_. If no **P’** is observed or one of the intensity scores of **P’** is higher than that of **P**, we discard **P**. If **P** is retained, we merge peak groups of masses ***m*** + ***nm***_***iso***_ for ***n*** = 0, 1, 2,…, sequentially, until there is no such peak group left. At the same time, we also recruit isotopologue peaks. Denote the mass of charge carrier by **q**. For each peak in **P** of charge **c**, we recruit peaks in the raw spectrum **S** having m/z of (***m*** + ***nm***_***iso***_+**cq**)/**c** to ***P*** for ***n*** = 0, 1, 2,…, until no such peak is found. We repeat the same procedure also for ***n*** = −1, −2,… This twoway scan ensures that the light (and less abundant) isotopologues are included in ***P***. During this merging and recruiting, any peak added to ***P*** at ***n*** =***i*** is assigned the (tentative) isotope index ***i***, which can be negative. After finishing the above procedure for **P**, all isotope indices are shifted so the minimum peak isotope index in **P** is 0 (still, tentative indices; see below). Each peak group now contains the peaks with different charges and different isotope indices that are likely to be from the same protein mass.

#### Peak group clustering for isotope envelopes (isotopically unresolved case)

In cases where isotopologues are not baseline resolved (in m/z direction) in raw profile spectrum, the corresponding peaks in the centroid spectrum (converted by mscovert in our case) often have inaccurate m/z positions. Furthermore, the peaks from distinct isotopologues are located so close to each other that they are within the mass tolerance. Thus, the above peak group clustering procedure results in the peak group **P** containing multiple occurance of the identical peaks with different isotope indices causing the scoring in the next step ineffective. To avoid this, we first retain only unique peaks in **P**. Then, for each unique peak, the isotope index **n** is assigned such that the distance between the peak mass and the mass of the **n**-th isotopologue (i.e., ***m*** + ***nm***_***iso***_) is minimized.

#### Filtration/scoring of peak groups and determination of the final isotope indices

To reduce false-positive masses, we filter out peak groups based on their per charge or per isotope intensity distribution. In this process, the determination of the final isotope indices (and thus the monoisotopic mass) is also carried out.

For a peak group, let the charge intensity for ***c*** be the summed intensity over the peaks of charge ***c*** in the peak group. We say a charge is intense if its charge intensity is higher than 25% of the max charge intensity. Let **m**_***i***_ be the maximum number of intense charges that continuously increment by **i**. For example, if charges 3, 5, 6, 7, 9, 10 are intense, **m**_**1**_ = 3 (charges 5, 6, 7) while **m**_**2**_ = 4 (charges 3, 5, 7, 9). A peak group is filtered out if **m**_**1**_ < max(3, **m**_**2**_,**m**_**3**_,…,**m**_**7**_). This filtration dramatically reduces remaining high harmonic artifacts.

Then the charge intensities are fitted to a Gaussian function using Caruana’s algorithm^22^, and if the cosine similarity between the charge intensities and the fitted Gaussian is less than 0.4 (user-specified parameter), we discard the peak group. Before the fitting, we find the smallest and largest charges such that their intensities exceed 2% of the maximum charge intensity. The charge intensities between the found charges (including zero charge intensities) are used as the input to Caruana’s algorithm. If Caruana’s algorithm yields a negative variance, the peak group is disregarded. Otherwise, the cosine similarity is calculated.

The isotope intensity for ***i*** (the summed intensity over the peaks of isotope index ***i*** in the peak group) is then used for further filtration. The isotope intensities are compared against the averagine distribution of the (tentative) monoisotopic mass of the peak group^23^. Let ***w*** be the isotope index of the maximum abundance in the averagine distribution. And denote the isotope intensity for ***i*** as ***I***_***i***_. As the isotope indices of the peaks might not be accurate, we give a large isotope index offset ***Δ*** from −***w*** to ***w*** and pick the offset ***Δ\**** that maximizes the cosine similarity between the isotope intensities ***I***_***i*** +***off***_ (for ***i*** =−***Δ***,…, 2***w***-***Δ***) and the avergine distribution. If the resulting maximum cosine similarity is less than 0.4 (user-specified), the peak group is filtered out. Otherwise, the isotope indices of all peaks in the peak group are shifted by ***Δ\**** (giving the final isotope indices). Monoisotopic mass is decided accordingly with the mass and the final isotope index of the most intense peak in the peak group.

#### Use of RT-adjacent spectral information

Similar to Xtract and ReSpect, FLASHDeconv can use information from spectra close in retention time. Given a spectrum ***S*** at RT ***rt*** and an RT window ***Δrt*** (user specified), we retain all the average masses of peak groups obtained from spectra at RTs between ***rt***- ***Δrt*** and ***rt***. When processing the spectrum ***S***, these masses are used to generate additional peak groups for the candidate peak groups; for these masses, the peak groups are formed and added even if there is no three consecutive signal peaks in ***S***. The next steps (i.e., assigning unique charges to peaks, isotope envelope clustering, and peak group filtration) are equally applied to these additional peak groups.

### Feature finding by mass trace detection

Once all spectra have been deconvolved as described above (and if there are multiple spectra), FLASHDeconv searches for features along the retention time. This idea goes back to the notion of ‘mass trace detection’ frequently used in quantitative mass spectrometry. We use the term feature here to describe all the signal arising from the same proteoform. Integrating this signal over retention time and summing it up across charge states/isotopes reduces the complexity of the data set and results in a feature primarily characterized by its retention time, mass, and intensity. Thus, feature detection is based on the notion of a mass trace in the deconvolved spectra.

We use a robust mass trace detection algorithm^19^ based on a Gaussian kernel density estimation of the local mass error and as implemented in OpenMS. For mass tolerance, twice the input tolerance to FLASHDeconv was used. The “re-estimate mass tolerance” function was activated. For each feature, the charge intensities and isotope intensities are defined similar to those for peak groups and are used for filtration of features (instead of peak groups) as above, but with higher cosine similarity score threshold (0.6, user-specified parameter).

### Cyto (Bovine Cytochrome C) and Fil (Filgrastim) data set acquisition

For the Cyto data set, Bovine Cytochrome C (P62894 UniProt accession number) was purchased from Sigma-Aldrich (Darmstadt, Germany). For the Fil data set, Filgrastim was kindly provided by CinnaGen Co. HPLC-grade H_2_O, acetonitrile (ACN) and formic acid (FA) were purchased from Merck (Darmstadt, Germany).

Each protein was dissolved with 0.1% FA in H_2_O to the concentration of 0.5 µg/µL. Reversed-phase liquid chromatography analysis was performed with an ultra-pressure liquid-chromatography (UPLC) system (ACQUITY, Waters, Manchester, UK). 0.1 % FA in H_2_O was used as mobile phase A and 0.1% FA in ACN was used as mobile phase B. 1 µL of sample were loaded onto a reversed-phase column (monolithic Proswift RP-4 Analytical 1 x 50 mm; Thermo Scientific, Bremen, Germany) and washed for 1 min with 2% mobile phase B. The proteins were eluted at a constant flow rate of 0.1 µL/min with a linear gradient increasing to 25% B in 5 min, 40% B in 5 min and 70% B in 3 min and hold in 70% B for 1 min. Column temperature was set to 30°C.

The eluting proteins were ionized via ESI and analyzed with Quadrupole-Orbitrap mass spectrometry (Q-Exactive, Thermo Scientific, Bremen, Germany) in a positive mode with 20.0 eV in-source CID. Full scan mass spectra were acquired at mass range 500 – 4,000 m/z, with a resolution of 70,000 and an AGC target of 3,000,000.

### PIP (Pierce Intact Protein Standard Mix) data set acquisition

Pierce Intact Protein Standard Mix containing six standard proteins namely Protein G, Protein AG, IGF-I LR3, Thioredoxin, Carbonic Anhydrase II, and Exo Klenow was purchased from Thermo Scientific (Bremen, Germany). The lyophilised powder of the mixture was reconstituted in 100 µL HPLC-grade water to a final concentration of 0.76 µg/µL. LC-MS analysis was performed as mentioned above with 0.1% FA water as mobile phase A, 0.1% FA acetonitrile as mobile phase B, and column temperature was set to 30°C. 1µL of sample was loaded onto a reversed-phase column (monolithic Proswift RP-4 Analytical 1 x 250 mm, Thermo Scientific, Bremen) and washed 2 min with 5% mobile phase B. The proteins were eluted at a constant flow rate of 0.2 µL/min with a linear gradient increasing to 25% B in 5 min, 45% B in the next 27 min. The MS data was obtained on a quadrupole-orbitrap mass spectrometer operated in positive mode with 20.0 eV in-source CID at a mass range of 500 to 3,000m/z, MS1 only at 17,500 resolution with 5 microscans and AGC target of 3,000,000.

### Tool parameters

#### FLASHDeconv

FLASHDeconv was used with default parameters for all data sets. By default, the mass range is set to 1-100 kDa and the charge range is 2-100. The isotope cosine threshold and charge intensity cosine threshold are set to 0.4 (spectrum level) and 0.6 (feature level). Mass tolerance is 10 ppm. The RT window is automatically set to 1% of total gradient time per data set.

#### Xtract

“Intact Protein Analysis” experiment type was used for all data sets. We used the processing method “Default SW Xtract” with the following modifications. “Output Mass Range” was set to 1-100 kDa for all data sets. In the case of “Charge Range”, 2-100 was used for the simple (Cyto, Fil, and PIP) data sets, but 2-50 for the myoblast data set, because Xtract often stopped working with 2-100 range for the myoblast data set. “Automatic Sliding Window Parameter Values” option was turned on for the simple data sets. But for the myoblast data set, the option was deactivated and “Target Avg Spectrum Width” was set to 3 minutes.

#### ReSpect

“Intact Protein Analysis” experiment type was used for all data sets. We used the processing method “Default SW ReSpect” with the following modifications. The mass ranges (both “Output Mass Range” and “Model Mass Range”) were set to 1-100 kDa and “Charge Range” to 2-100 for all data sets. “Target Mass” was set to 50 kDa. “Automatic Sliding Window Parameter Values” option was turned on for the simple data sets. But for the myoblast data set, the option was deactivated and “Target Avg Spectrum Width” was set to 3 minutes.

#### Promex

For all data sets, the mass range was set to 1-100 kDa, and the charge range was set to 2-60 (the maximum charge for Promex is 60).

### Mass artifact and isotopologue detection in a feature list

Given a set of features **F**, to test if a feature ***f*** in **F** with mass ***m*** is a mass artifact or one of isotopologues, we first collected all features in ***F*** such that the RT overlap between the feature and ***f*** exceeds 60% of the RT span of ***f***. Secondly, the features having lower intensity than ***f*** are filtered out. Denote the remaining feature set by ***F***_***f***_. To determine, for example, if ***f*** is a low harmonic artifact, we tested if there exists a feature in ***F***_***f***_ that has mass of ***cm*** for ***c*** = 2,…, 100 (within 10 ppm tolerance) allowing up to 10 isotope error. If such a feature exists, we declared ***f*** as a low harmonic artifact feature. More formally, ***f*** is a low harmonic artifact if there exists a feature in ***F***_***f***_ whose mass equals ***c*** (***m*** + ***km_iso_***) within 10 ppm tolerance for ***c*** =2,…,100 and ***k*** = −10,…, 10, where ***m_iso_*** denotes the mass difference between 13C and 12C. For high harmonic artifacts, the mass term (***m*** + ***km_iso_***)/***c*** was used in the place of ***c*** (***m*** + ***km_iso_***).

Charge off by one artifact features^11^ occur when a charge ***c*** +1 or ***c***-1 is assigned to a charge ***c*** peak. They mainly arise when the charge assignment is based on the peak m/z distance between isotopologues because the measurement of the distance for highly charged peaks is often inaccurate, not to mention the isotopically unresolved peaks. But even when the charge assignment is based on the charge pattern, the artifact can be generated if two peaks with consecutive charge states are close to each other (e.g, below 4 Th), due to peak m/z variations. Thus, to detect the charge off by one artifacts, we chose the charge states ***c\**** (between 2 and 100) such that ***m*** /***c\****− **m**/(***c\**** +1)<4. For all such ***c\****, we tested if features having masses (***m*** +***km_iso_***)*(***c\**** +1)/***c\**** or (***m*** +***km_iso_***)*(***c\****−1)/***c\**** are present in ***F***_***f***_ (10 ppm mass tolerance allowed) for ***k*** = −10,…, 10. If present, we declared ***f*** to be a charge off by one artifact.

Lastly, to see if ***f*** has other features corresponding to its isotopologues, we simply checked if features having masses ***m*** − ***m_iso_*** or ***m*** + ***m_iso_***, within 10 ppm tolerance, are present in ***F***_***f***_. If present, ***f*** is considered to be one of the isotopologues. We repeat the above procedure for each feature in the list ***F***.

### Matching feature masses against protein masses identified by bottom-up searches

Feature masses from complex data sets were matched to the protein masses identified by BU search, from the same myoblast sample. To collect the protein masses to be matched, we took the UniProt (downloaded in April, 2019) entries identified by BU search in L. V. Schaffer et al., 2018^14^. The (unmodified) protein masses were then calculated using their amino acid sequences. For the matches with modifications, 18 relatively abundant modification types found by PTMCurator^24^ and two protein N-terminal modifications (Met-loss and Met-loss+Acetyl) were considered (see Supplementary Table 2). We performed two matchings: one with zero modification and the other with up to one modification per protein.

Matching between a feature mass and the set of all theoretical proteoform masses (subject to the above modification criteria) was done by using the multimap data structure. For each possible proteoform mass, the multimap takes the mass as key and the proteoform(s) having the mass as value. A feature is said to be matched to a proteoform if the key (i.e., mass) of the proteoform is within 10 ppm tolerance from the feature mass, allowing one isotope error.

## Supplementary information

**Supplementary Figure 1. Analogue of Figure 1b-e for the simple data sets (Cyto, Fil, and PIP) technical replicate 1.**

**Supplementary Figure 2. Analogue of Figure 1b-e for the simple data sets (Cyto, Fil, and PIP) technical replicate 2.**

**Supplementary Figure 3. Venn diagram for the overlap of the rounded monoisotopic masses between tools in low mass region for the myoblast data set.**

**Supplementary Data 1. The outputs from FLASHDeconv, Xtract, ReSpect, and Promex for the simple data sets (Cyto, Fil, and PIP).**

**Supplementary Data 2. The outputs from FLASHDeconv, Xtract, ReSpect, and Promex for the simple data sets (Cyto, Fil, and PIP) technical replicate 1.**

**Supplementary Data 3. The outputs from FLASHDeconv, Xtract, ReSpect, and Promex for the simple data sets (Cyto, Fil, and PIP) technical replicate 2.**

**Supplementary Data 4. The outputs from FLASHDeconv, Xtract, ReSpect, and Promex for the myoblast data set.**

**Supplementary Table 1. Features that matched to the protein masses identified by bottom-up (BU) search for the myoblast data set.**

**Supplementary Table 2. The list of modifications considered in the feature-BU identified protein mass matching.**

