## Supplementary materials for "FLASHDeconv: Ultrafast, high-quality feature deconvolution for top-down proteomics"

### Supplementary materials for FLASHDeconv: Ultra-fast, high-quality feature deconvolution for top-down proteomics

**Figure 1.**

**a.**

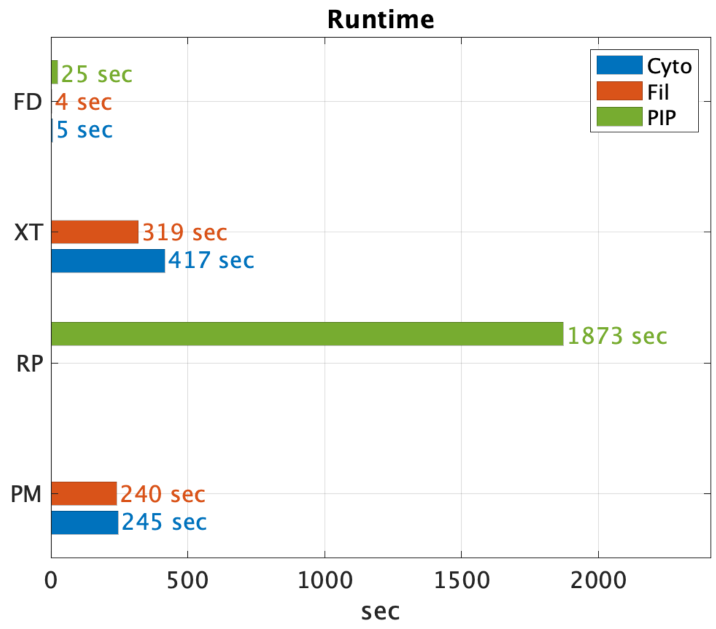
Runtime
Cyto
Fil
PIP
FD
XT
RP
PM
25 sec
4 sec
5 sec
319 sec
417 sec
1873 sec
240 sec
245 sec
0
500
1000
1500
2000
sec

**b.**

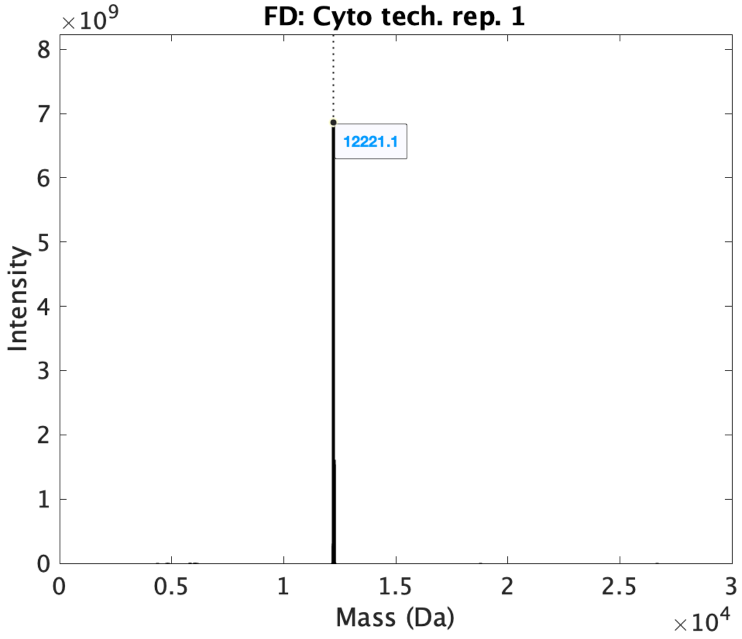
FD: Cyto tech. rep. 1
×10^9
8
7
6
5
4
3
2
1
0
Intensity
12221.1
0
0.5
1
1.5
2
2.5
3
Mass (Da)
×10^4

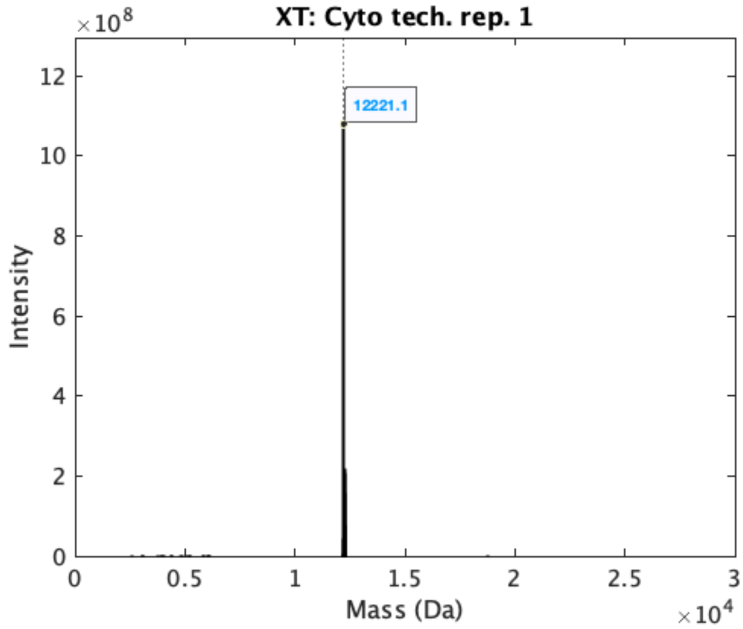
XT: Cyto tech. rep. 1
×10^8
12
10
8
6
4
2
0
Intensity
12221.1
0
0.5
1
1.5
2
2.5
3
Mass (Da)
×10^4

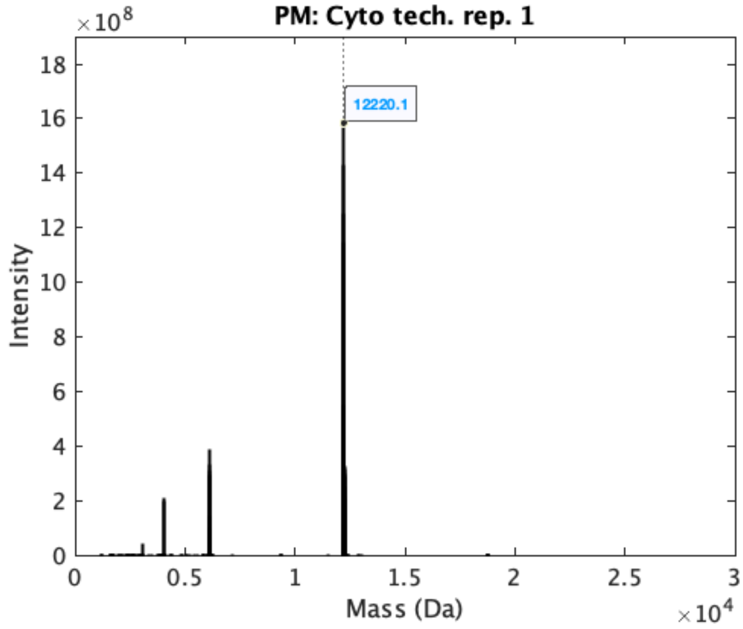
PM: Cyto tech. rep. 1
×10^8
18
16
14
12
10
8
6
4
2
0
Intensity
12220.1
0
0.5
1
1.5
2
2.5
3
Mass (Da)
×10^4

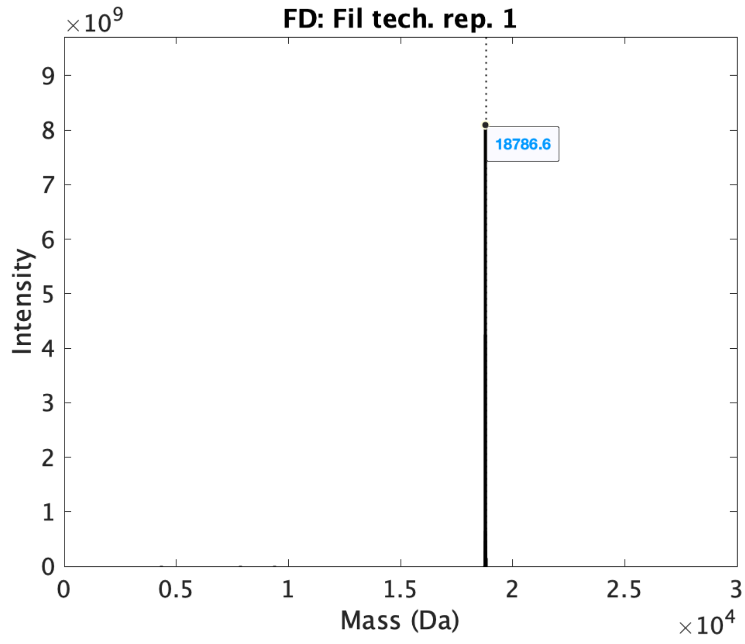
FD: Fil tech. rep. 1
×10^9
9
8
7
6
5
4
3
2
1
0
Intensity
18786.6
0
0.5
1
1.5
2
2.5
3
Mass (Da)
×10^4

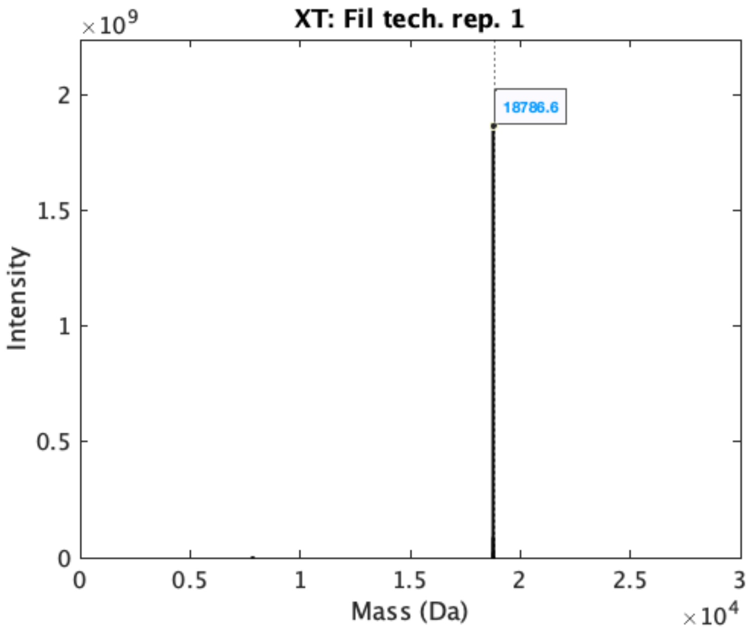
XT: Fil tech. rep. 1
×10^9
2
1.5
1
0.5
0
Intensity
18786.6
0
0.5
1
1.5
2
2.5
3
Mass (Da)
×10^4

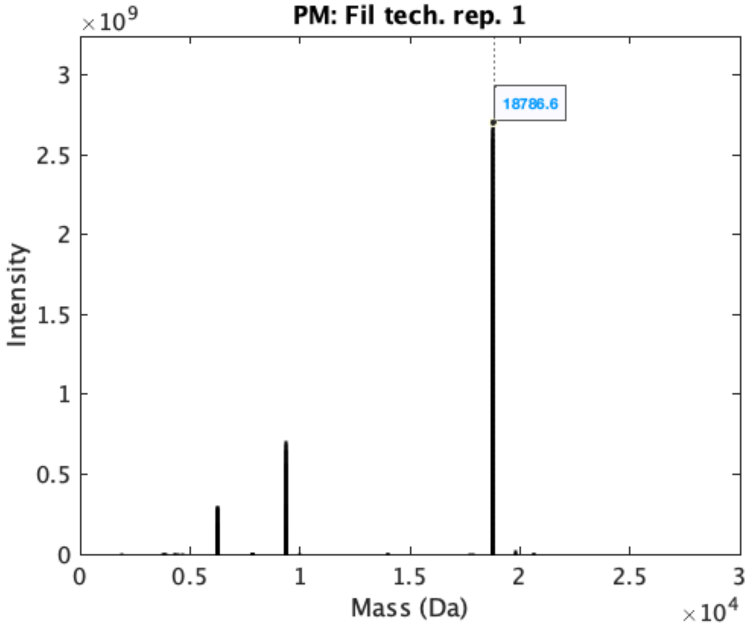
PM: Fil tech. rep. 1
×10^9
3
2.5
2
1.5
1
0.5
0
Intensity
18786.6
0
0.5
1
1.5
2
2.5
3
Mass (Da)
×10^4

**c.**

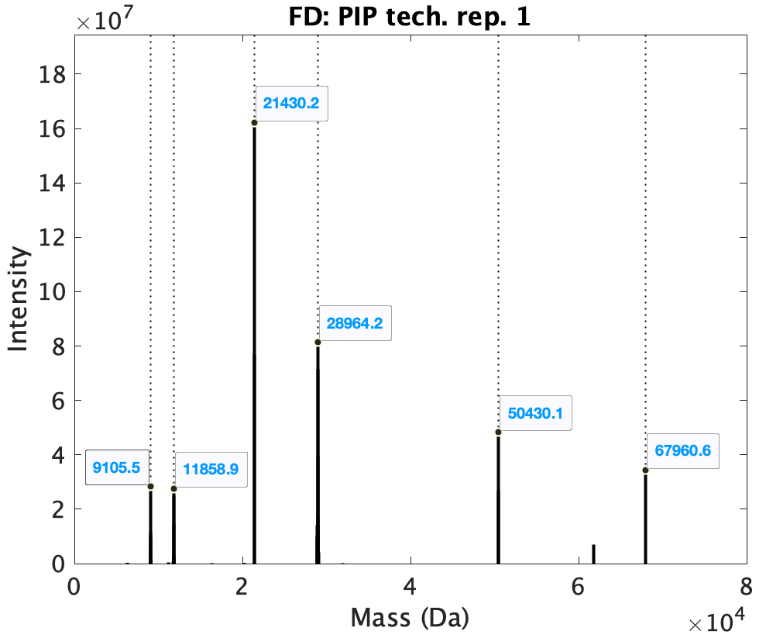
FD: PIP tech. rep. 1
×10^7
Intensity
Mass (Da)
×10^4
9105.5
11858.9
21430.2
28964.2
50430.1
67960.6

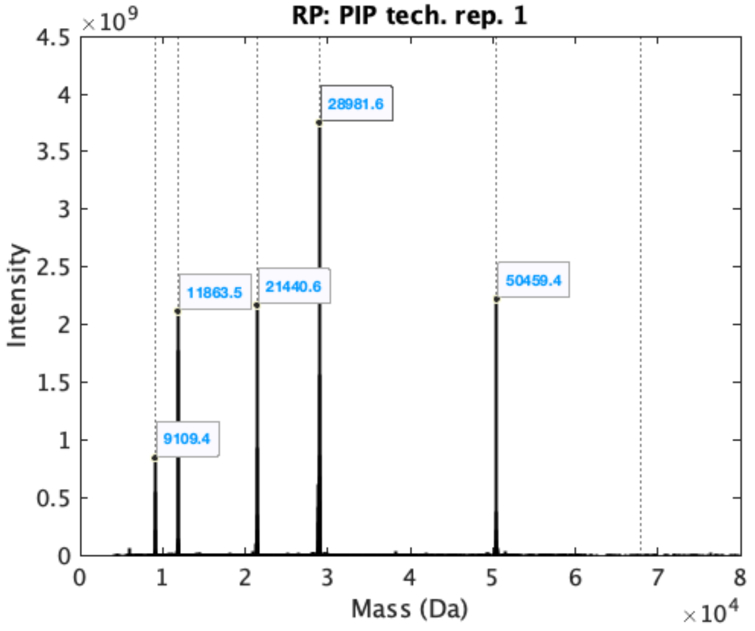
RP: PIP tech. rep. 1
×10^9
Intensity
Mass (Da)
×10^4
9109.4
11863.5
21440.6
28981.6
50459.4

**d.**

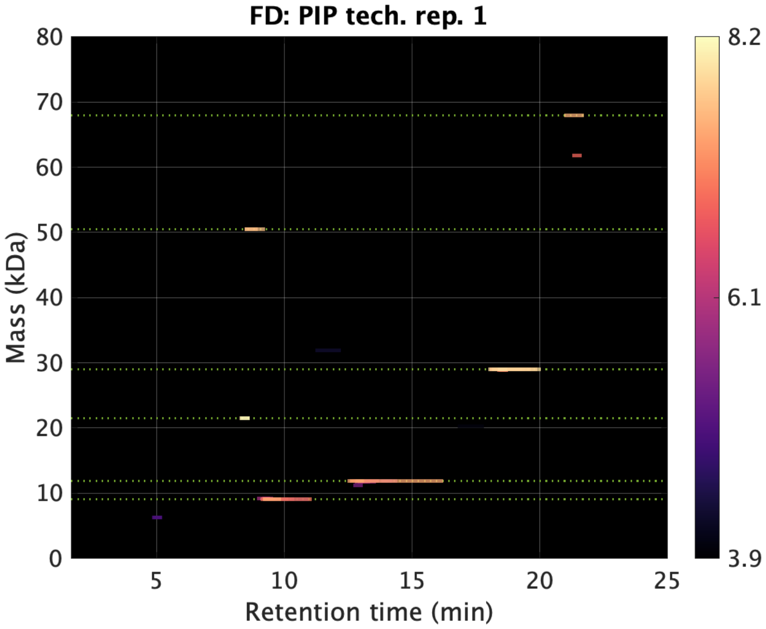
FD: PIP tech. rep. 1
Mass (kDa)
Retention time (min)
0
10
20
30
40
50
60
70
80
5
10
15
20
25
8.2
6.1
3.9

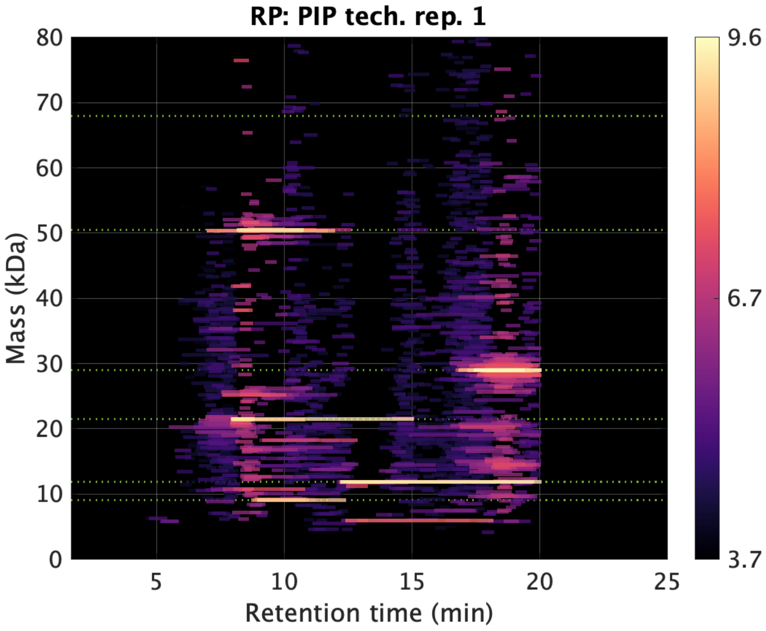
RP: PIP tech. rep. 1
Mass (kDa)
Retention time (min)
0
10
20
30
40
50
60
70
80
5
10
15
20
25
9.6
6.7
3.7

**Figure 1. Analogue of Figure 1b-e for the simple data sets (Cyto, Fil, and PIP) technical replicate 1. a.** Runtime comparison between tools. **b.** Deconvoluted spectra from the Cyto and Fil (isotopically resolved) data sets. **c.** Deconvoluted spectra from the PIP (isotopically unresolved) data set. **d.** Features maps from the PIP data set.

**Figure 2.**

**a.**

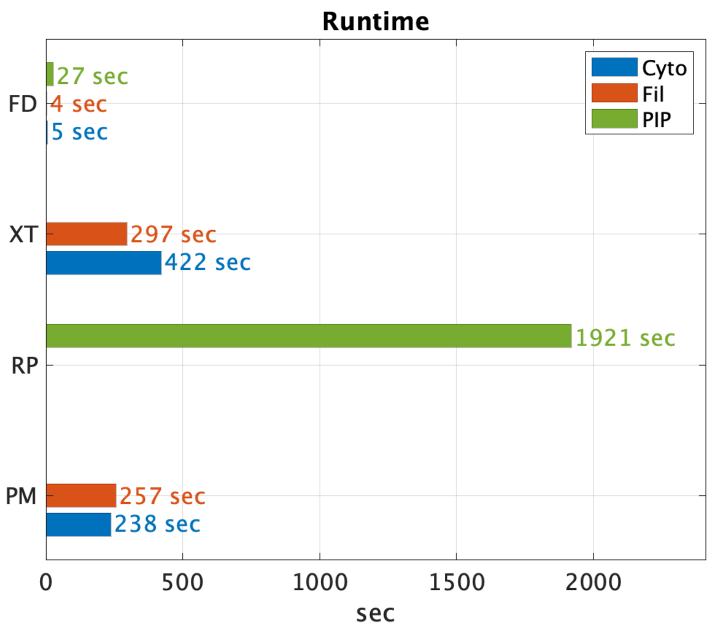

Runtime
Cyto
Fil
PIP
FD
27 sec
4 sec
5 sec
XT
297 sec
422 sec
RP
1921 sec
PM
257 sec
238 sec
0
500
1000
1500
2000
sec


**b.**

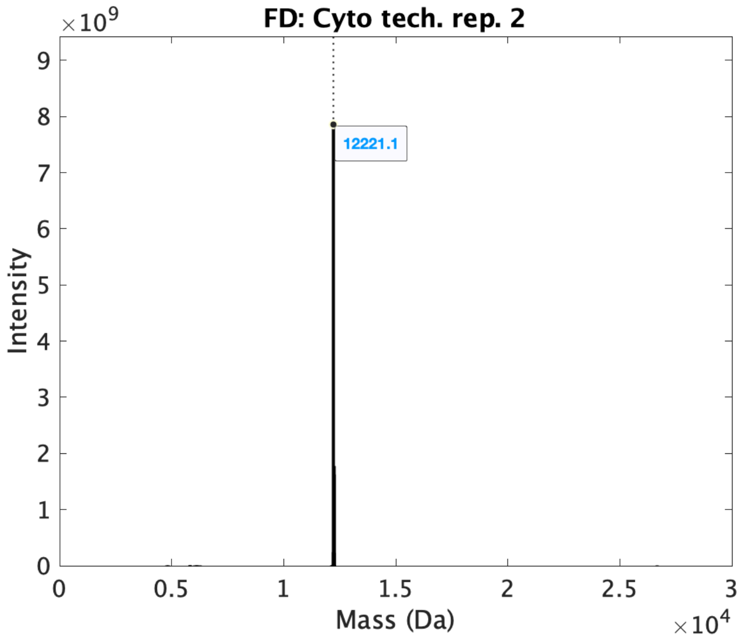

FD: Cyto tech. rep. 2
×10^9
Intensity
12221.1
Mass (Da)
×10^4

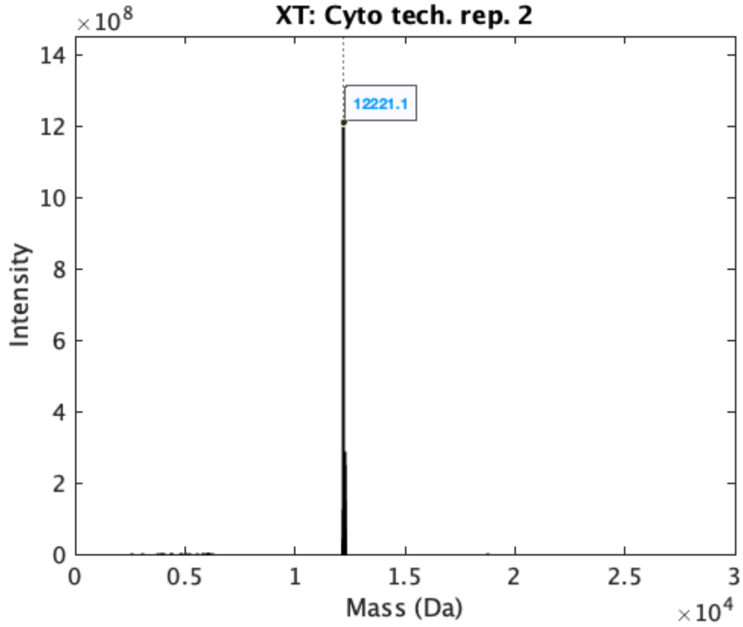
XT: Cyto tech. rep. 2
×10^8
14
12
10
8
6
4
2
0
Intensity
12221.1
0
0.5
1
1.5
2
2.5
3
Mass (Da)
×10^4

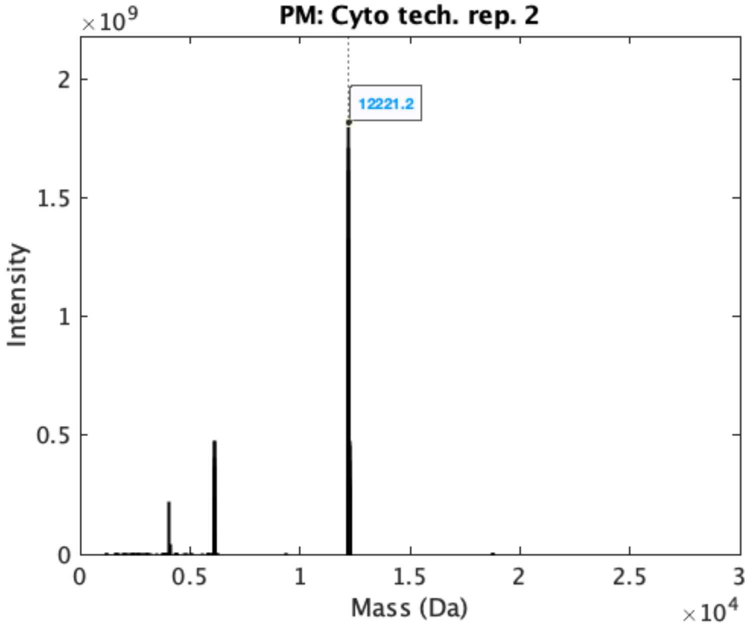
PM: Cyto tech. rep. 2
×10^9
2
1.5
1
0.5
0
Intensity
12221.2
0
0.5
1
1.5
2
2.5
3
Mass (Da)
×10^4

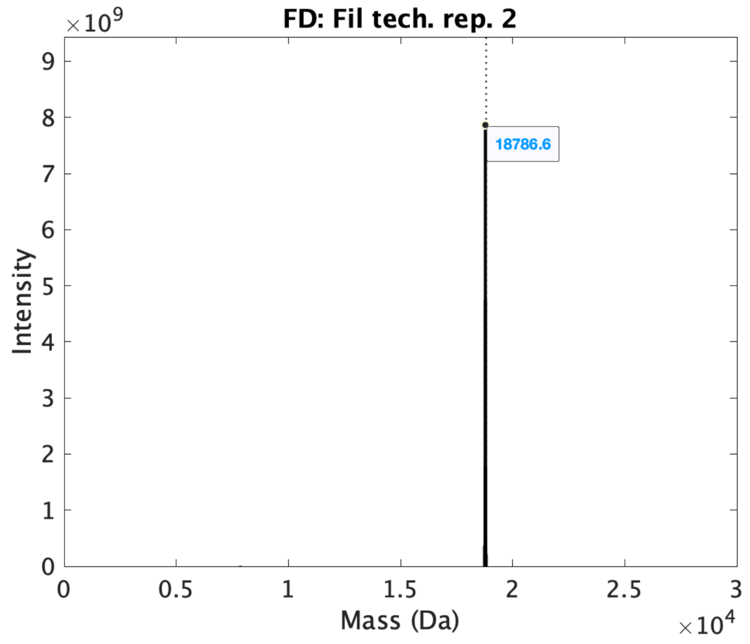
FD: Fil tech. rep. 2
×10^9
Intensity
Mass (Da)
×10^4
18786.6

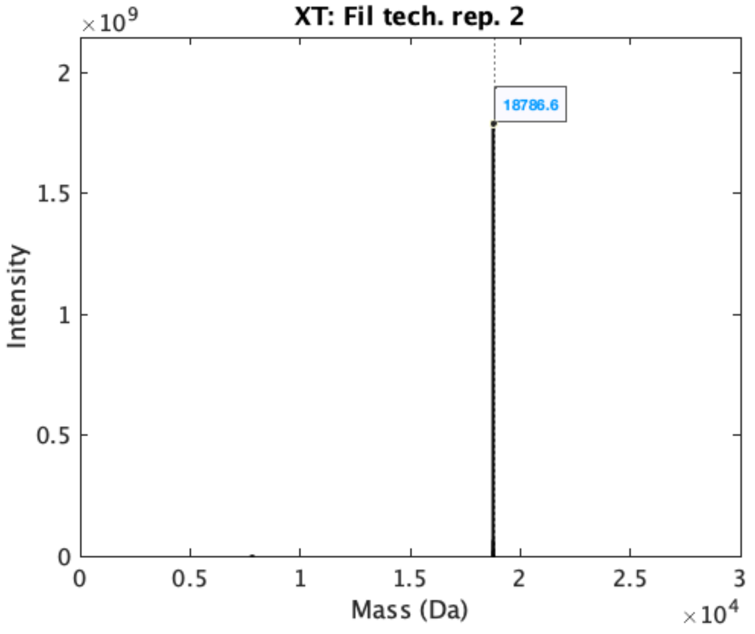
XT: Fil tech. rep. 2
×10^9
Intensity
Mass (Da)
×10^4
18786.6

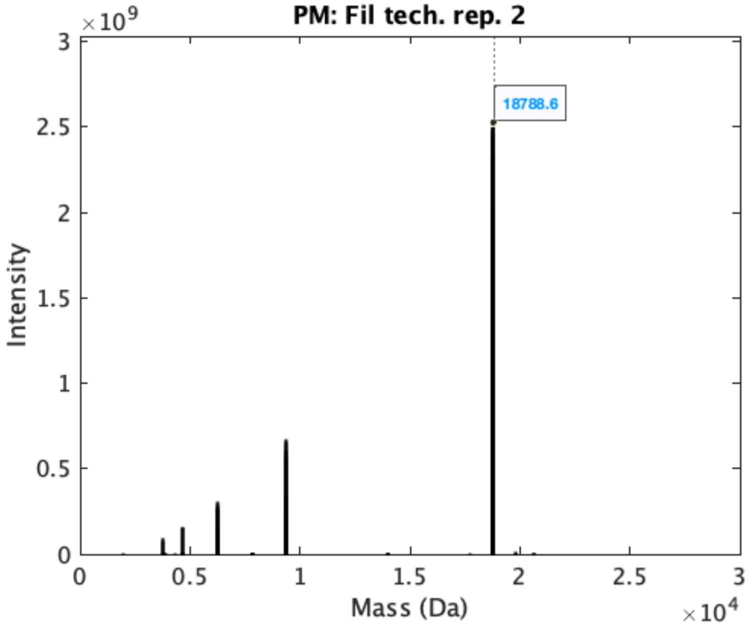
PM: Fil tech. rep. 2
×10^9
18788.6
Intensity
Mass (Da)
×10^4
0
0.5
1
1.5
2
2.5
3

**c.**

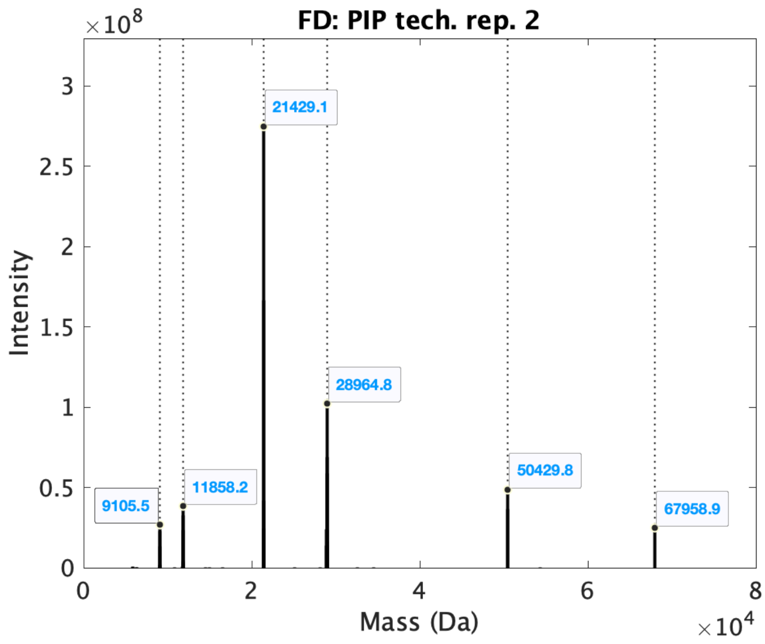
FD: PIP tech. rep. 2
×10^8
Intensity
Mass (Da)
×10^4
9105.5
11858.2
21429.1
28964.8
50429.8
67958.9

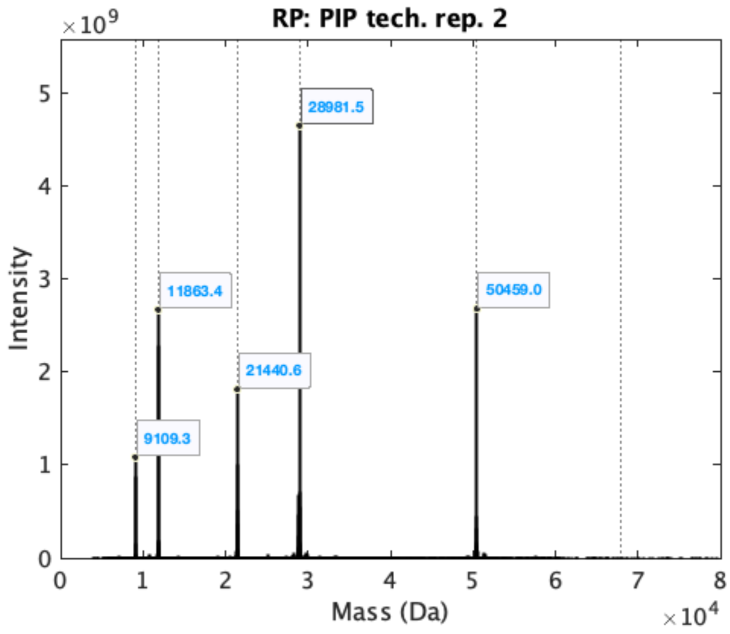
RP: PIP tech. rep. 2
×10^9
Intensity
Mass (Da)
×10^4
9109.3
11863.4
21440.6
28981.5
50459.0

d.

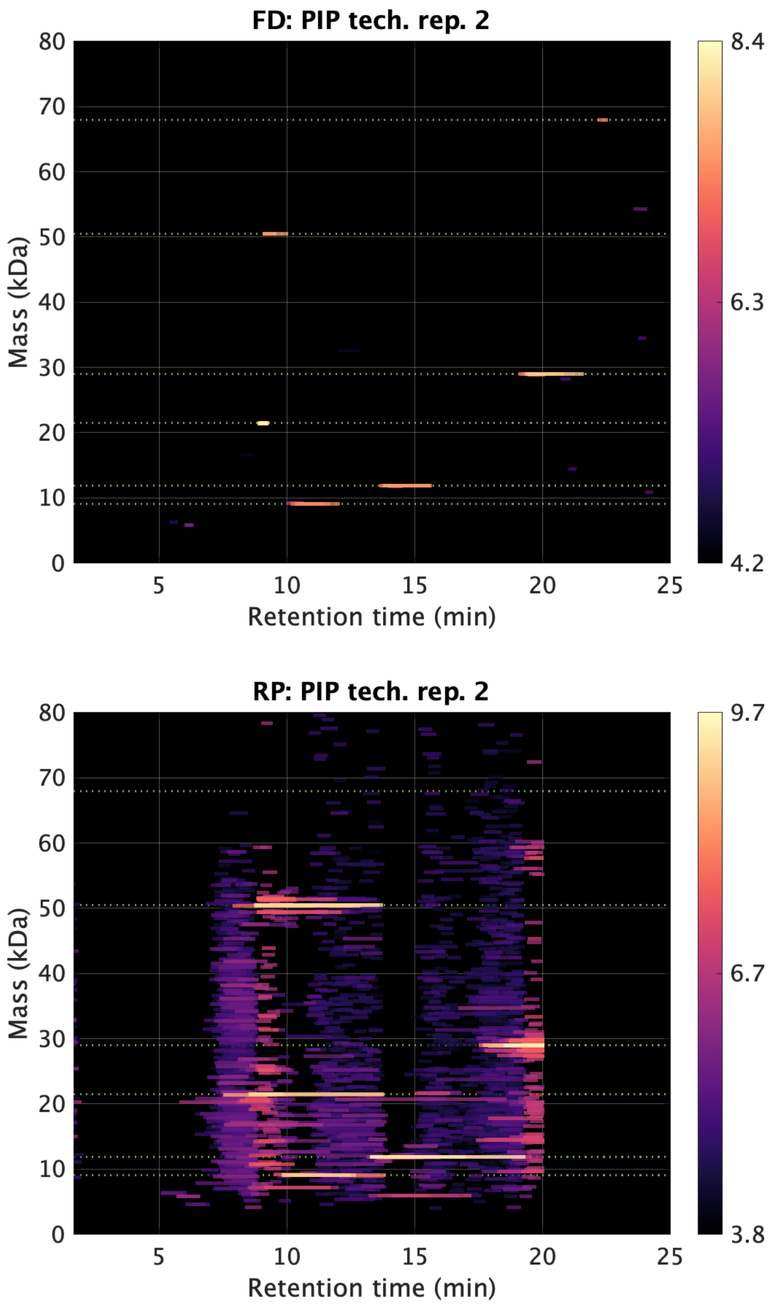


**Figure 2.** **Analogue of Figure 1b-e for the simple data sets (Cyto, Fil, and PIP) technical replicate 2.**

**Figure 3.**

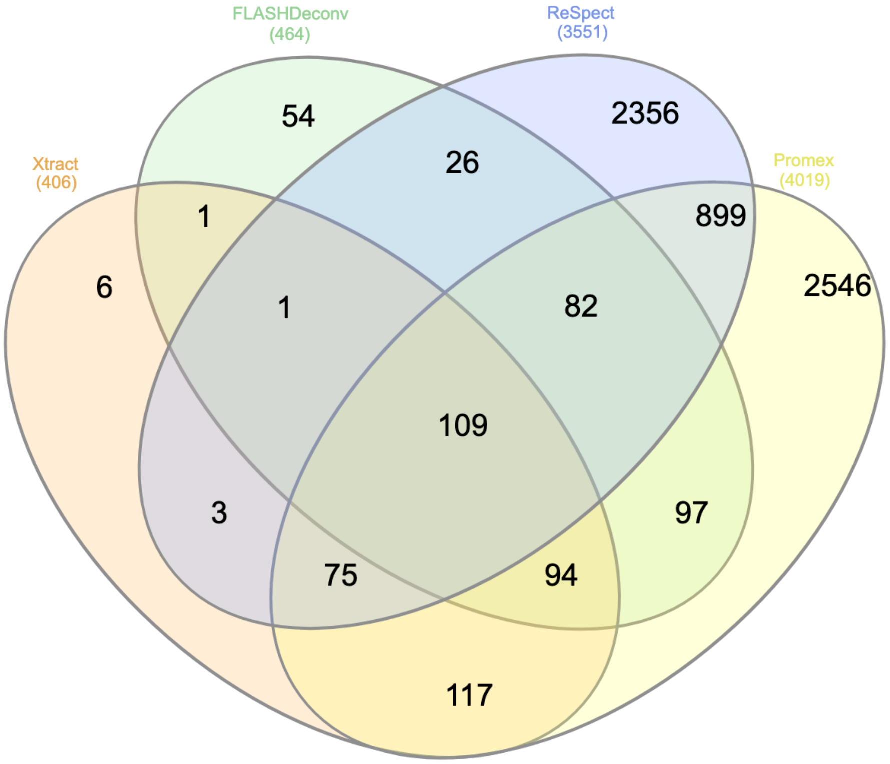


**Figure 3. Venn diagram for the overlap of the rounded monoisotopic masses between tools in low mass region for the myoblast data set.** For each tool, the features of mass less than 20 kDa and of RT 40-80 min were collected. For RP, monoisotopic masses were calculated from average masses using averagine model[1]. About 88% of FD masses overlapped with the other tools. The venn diagram was drawn using InteractiVenn[2].

| Tool | # features | Unmod | 1 PTM |
|---|---|---|---|
| **FD** | 6,969 | 579 (8%) | 5,176 (74%) |
| **XT** | 594 | 13 (2%) | 176 (30%) |
| **RP** | 20,258 | 1,208 (6%) | 11,870 (59%) |
| **PM** | 6,653 | 155 (2%) | 1,800 (27%) |

**Table 1. Features that matched to the protein masses identified by bottom-up (BU) search for the myoblast data set.** This table compares the number/portion of features that are matched to the protein masses from BU searches for myoblast sample (also shown in Fig. 2f). The second column gives the total number of reported features per tool. The third and fourth columns show the numbers (and portions) of the matched features when no PTM and 1 PTM were allowed, respectively (see online methods for details).

| PSI-MS name | Monoisotopic mass |
|---|---|
| Phospho | 79.966331 |
| Acetyl | 42.010565 |
| Amidated | -0.984016 |
| Oxidation | 15.994915 |
| Methyl | 14.015650 |
| LRGG | 383.228103 |
| Glu->pyro-Glu | -18.010565 |
| Gln->pyro-Glu | -17.026549 |
| Carboxy | 43.989829 |
| Palmitoyl | 238.229666 |
| Myristoyl | 210.198366 |
| ADP-Ribosyl | 541.061110 |
| Farnesyl | 204.187801 |
| Nitrosyl | 28.990164 |
| GeranylGeranyl | 272.250401 |
| Formyl | 27.994915 |
| Deamidated | 0.984016 |
| Sulfo+amino (Interim name) | 94.967714 |
| Met-loss | -131.040485 |
| Met-loss+Acetyl | -89.029920 |

**Table 2. The list of modifications considered in the feature-BU identified protein mass matching.**

#### Supplementary materials references

1. Senko, M. W., et al. *J Am Soc Mass Spectrom* **6**, 229-233 (1995).

2. Heberle, H., et al. *Bmc Bioinformatics* **16** (2015).
